# www.Malaria.tools - comparative genomic and transcriptomic database for *Plasmodium* species

**DOI:** 10.1101/639179

**Authors:** Qiao Wen Tan, Marek Mutwil

## Abstract

Malaria is a tropical parasitic disease caused by the *Plasmodium* genus, which resulted in an estimated 219 million cases of malaria and 435,000 malaria-related deaths in 2017. Despite the availability of the *P. falciparum* genome since 2002, almost 50% of the genes remain unannotated. To remedy this paucity of functional information, we used transcriptomic data to build gene co-expression networks for two *Plasmodium* species (*P. falciparum and P. berghei*), and included genomic data of four other *Plasmodium* species, *P. yoleii, P. knowlesi, P. vivax* and *P. cynomolgi*, as well as two non-*Plasmodium* species from the Apicomplexa, *Toxoplasma gondii* and *Theileria parva.* The database is preloaded with tools that allow the identification and cross-species comparison of co-expressed gene neighborhoods, clusters, and life stage-specific expression, thus providing sophisticated tools to predict gene function. Moreover, we exemplify how the tools can be used to easily identify genes relevant for pathogenicity and various life stages of the malaria parasite. The database is freely available at www.malaria.tools.

## INTRODUCTION

Malaria is a widespread infectious disease transmitted by the *Anopheles* mosquito which caused an estimated 219 million cases and 435,000 malaria-related deaths in the year 2017 (1). It is caused by various *Plasmodium* species with *P. falciparum* and *P. vivax* being the two most widespread and deadly for humans (2). Over the years, varying degrees of resistance have emerged in *Plasmodium* against all drugs used for treating malaria. Hence, it is a race against time to find new treatments, and this requires an understanding of malaria biology, and more specifically, the characterisation of genes and their functions in order to develop drugs that target genes responsible for pathogenicity.

Despite the availability of genomes of the various *Plasmodium* parasites for already more than a decade, many of the genes have unknown functions. These genes are often parasite-specific (3), thus making conventional gene function prediction methods which are based on homology to genes from other organisms less efficient. To this end, alternative methods for gene function prediction such as gene co-expression networks have been developed. The rationale behind using gene co-expression networks stems from the observation that functionally related genes have similar gene expression profiles (4). Thus, by identifying clusters of genes that have highly similar expression profiles, we attain groups of functionally related genes where the function of uncharacterised genes can be inferred from its neighbours.

We have built the database Malaria.tools (www.malaria.tools) based on the CoNekT database framework (5) using publicly available RNA sequencing data of two model *Plasmodium* species. This database calculates gene co-expression networks from gene expression data obtained from more than 800 experiments and contains multiple tools to predict gene function from co-expression network neighbours, co-expressed clusters and specific expression profiles. Additionally, an in-built phylogenetic tree function combined with gene expression comparison allows *Plasmodium* researchers to identify orthologs of *P. falciparum* genes. Malaria.tools provides the malaria research community a comprehensive and highly valuable resource for an efficient characterisation of genes based on their expression profile, and will aid in the identification of potential drug targets *in silico* in the race for new antimalarial drugs.

## MATERIALS AND METHODS

RNA sequencing experiments for *P. falciparum* and *P. berghei* were downloaded as fastq files from European Nucleotide Archive (ENA) (6) via aspera v3.8.1.160447. For paired end experiments, only the file containing the first read, designated with “_1” was downloaded. To remove possible RNA contaminants from the parasite’s hosts, the reads were mapped first against human and mouse for *P. falciparum* and *P. berghei*, respectively. The unmapped reads were then mapped against the mosquito vector. Human mosquito vector was used for both *Plasmodium* species as the CDS of the mouse mosquito vector was unavailable. Unmapped reads were mapped against the respective *Plasmodium* species, and experiments with at least 1 million reads and 50% of reads mapped to the *Plasmodium* species were used to construct the database.

The mapping was done using kallisto v0.44.0 (7). Kallisto index files were generated for *Homo sapiens* (GCF_000001405.38)*, Anopheles gambiae* (GCF_000005575.2)*, P. falciparum* 3D7 and *P. berghei* ANKA CDS sequences with default parameters. Both paired and single libraries were treated as single end libraries for mapping using kallisto quant for single end library with estimated fragment length of 200bp, estimated standard deviation of 20 and pseudobam option. Unmapped reads from the output BAM files were written into a new fastq file using samtools v1.9-52-g651bf14 (8).

In total, 206 experiments from *P. berghei* (Figure S1A) and 620 experiments from *P. falciparum* (Figure S1B) were included in database and annotated based on the information available from NCBI Sequence Read Archive, ENA and literature (Supplementary Table S3 (*P. berghei)* and Supplementary Table S4 (*P. falciparum*)). The Transcripts Per Kilobase Million (TPM) values from kallisto output of the selected experiments were represented as an expression matrix where genes are arranged in rows and experiments in columns. The expression matrices are available in Table S1 for *P. berghei* and Table S2 for *P. falciparum*. Highest Reciprocal Rank (HRR) co-expression networks were then constructed (9).

CDS sequences, description and associated GO terms for the 8 species in the database were obtained from the various sources described in Table 1. Superfamily and Pfam domain annotations from interproscan v5.32-71.0 (10) were used as sequence description when descriptions were not available. Orthologous groups of genes (Supplementary Table S5) and phylogenetic trees were obtained from Orthofinder v1.1.8 (11) using BLAST for sequence similarity inference with default settings. The database was based on the CoNekt database framework (5) with default settings, where Heuristic Cluster Chiseling Algorithm (HCCA) (9) cluster size was limited to 100 genes. Experiments involving wild type *Plasmodium* were further binned according to its various life stages (ring, trophozoite, schizont, male gametocyte and female gametocyte) and used to calculate tissue specificity. Nucleotide and protein blast databases for blastn and blastp were created using makeblastdb v2.9.0+ (12).

**Table 1.** Source of data for species in database.

| Species | RNA-seq samples | CDS | CDS description | GO annotation |
| --- | --- | --- | --- | --- |
| *Theilera parva* Muguga | N/A | NCBI Ref-seq | NCBI Ref-seq | Interproscan |
| *Toxoplasma gondii* ME49 | N/A | NCBI Ref-seq | NCBI Ref-seq | Interproscan |
| *Plasmodium berghei* ANKA | 206 | PlasmoDB | GeneDB | PlasmoDB |
| *Plasmodium cynomolgi* B | N/A | PlasmoDB | Interproscan | Interproscan |
| *Plasmodium falciparum* 3D7 | 620 | PlasmoDB | GeneDB | PlasmoDB |
| *Plasmodium knowlesi* Malayan Pk1A | N/A | PlasmoDB | Interproscan | Interproscan |
| *Plasmodium vivax* P01 | N/A | PlasmoDB | GeneDB | Interproscan |
| *Plasmodium yoelii* YM | N/A | PlasmoDB | Interproscan | Interproscan |

## RESULTS

Malaria.tools offers a wide selection of tools to query the database. For example, the user can find the genes of interest by using BLAST, gene IDs (e.g., *PF3D7_1223100*) and keywords (e.g., rhoptry). Genes that work together in a specific biological process or contain a particular domain can be identified by querying the database with Gene Ontology terms (e.g., GO:0009405) or a Pfam domain (e.g., VSA_Rifin), respectively. The database offers multiple comparative genomic and transcriptomic tools that allow the user to view and compare expression profiles within and across species, and to investigate the phylogenetic and expression relationships of gene families. A full description of the features is found at https://malaria.sbs.ntu.edu.sg/features. To exemplify some of the features of malaria.tools, we provide three analyses showing typical case studies.

### Identification of a co-expression neighborhood important for erythrocyte invasion

A co-expression neighbourhood consists of a gene of interest and its co-expressed genes (neighbors) calculated based on Highest Reciprocal Rank (HRR). To identify co-expression neighborhoods of interest to malaria researchers, we calculated which genes are network neighbors to already functionally characterized genes. We identified gene *PF3D7_1223100*, which was co-expressed with 91 other genes of which 48 (53%) are annotated with specific GO terms indicating that experimental evidence for the gene function already exists (e.g. evidence codes EXP, IDA, IPI). The high percentage of functionally characterized genes in this neighborhood indicates that the corresponding biological process has received special attention from the malaria researchers, likely due to its involvement in pathogenicity.

To gain insight into the function of this neighborhood, we first studied the expression profile of *PF3D7_1223100* (https://malaria.sbs.ntu.edu.sg/sequence/view/16011). The expression profiles in malaria.tools are available in a detailed format that showcase expression in all annotated samples (Figure 1A), and as an average expression in the major life stages of malaria (Figure 1B, https://malaria.sbs.ntu.edu.sg/profile/view/8699). In both expression profiles, we observed that *PF3D7_1223100* and its co-expressed genes are expressed in all major life stages with particularly high expression in the schizont stage (Figure 1A,B). Next, we retrieved publications on the functional characterization of the genes and we found that the highly studied genes are clearly associated with functions important for erythrocyte invasion. The genes can be classified into 3 major groups relating to motility, cytoadherence and erythrocyte invasion. Genes relating to the glideosome complex [*PF3D7_0918000* (GAP50) (13), *PF3D7_1323700* (GAPM1) (14)], the inner membrane complex (15) [*PF3D7_0109000* (PHIL1) and *PF3D7_1003600* (IMC1c)] and cytoskeleton (16, 17) [*PF3D7_1251200* (coronin), *PF3D7_0932200* (Profilin) and *PF3D7_1246200* (Actin 1)] are essential for the parasite to move towards a new red blood cell through gliding motility. Upon reaching the red blood cell, merozoite surface proteins such as *PF3D7_1035400* (MSP3), *PF3D7_1335100* (MSP7), *PF3D7_1035500* (MSP6) (18) and *PF3D7_1035700* (DBLMSP) facilitate the binding of the parasite to the red blood cell. Finally, erythrocyte invasion is enabled by various genes such as enzymes [*PF3D7_0507500* (SUB1) (19), *PF3D7_1136500* (casein kinase 1) (20) and *PF3D7_0404700* (DPAP3) (21), signalling mediators (22) [*PF3D7_0934800* (PKAc), *PF3D7_1223100* (PKAr)], rhoptry proteins (23, 24) (*PF3D7_0929400* (RhopH2), *PF3D7_0905400* (RhopH3), *PF3D7_1410400* (RAP1), *PF3D7_0414900* (ARO), *PF3D7_1017100* (RON12), *PF3D7_0501600* (RAP2) and *PF3D7_0817700* (RON5)] and others [*PF3D7_0423800* (CyRPA) (25), *PF3D7_0935800* (CLAG9) (26), *PF3D7_0612700* (P12), *PF3D7_0404900* (P41) (27)]. The schizont is a non-infective life stage during the erythrocytic cycle. However, a mature schizont contains multiple merozoites, which upon rupture of the schizont moves and invades fresh erythrocytes. In the co-expression neighbourhood of *PF3D7_1223100*, we observe an upregulation of merozoite and erythrocyte invasion-related genes during the schizont stage. In conclusion, the remaining 47% of genes in this cluster that are not yet functionally characterized are prime candidates for further studies on parasite motility, cytoadherence and erythrocyte invasion.

**Figure 1.**
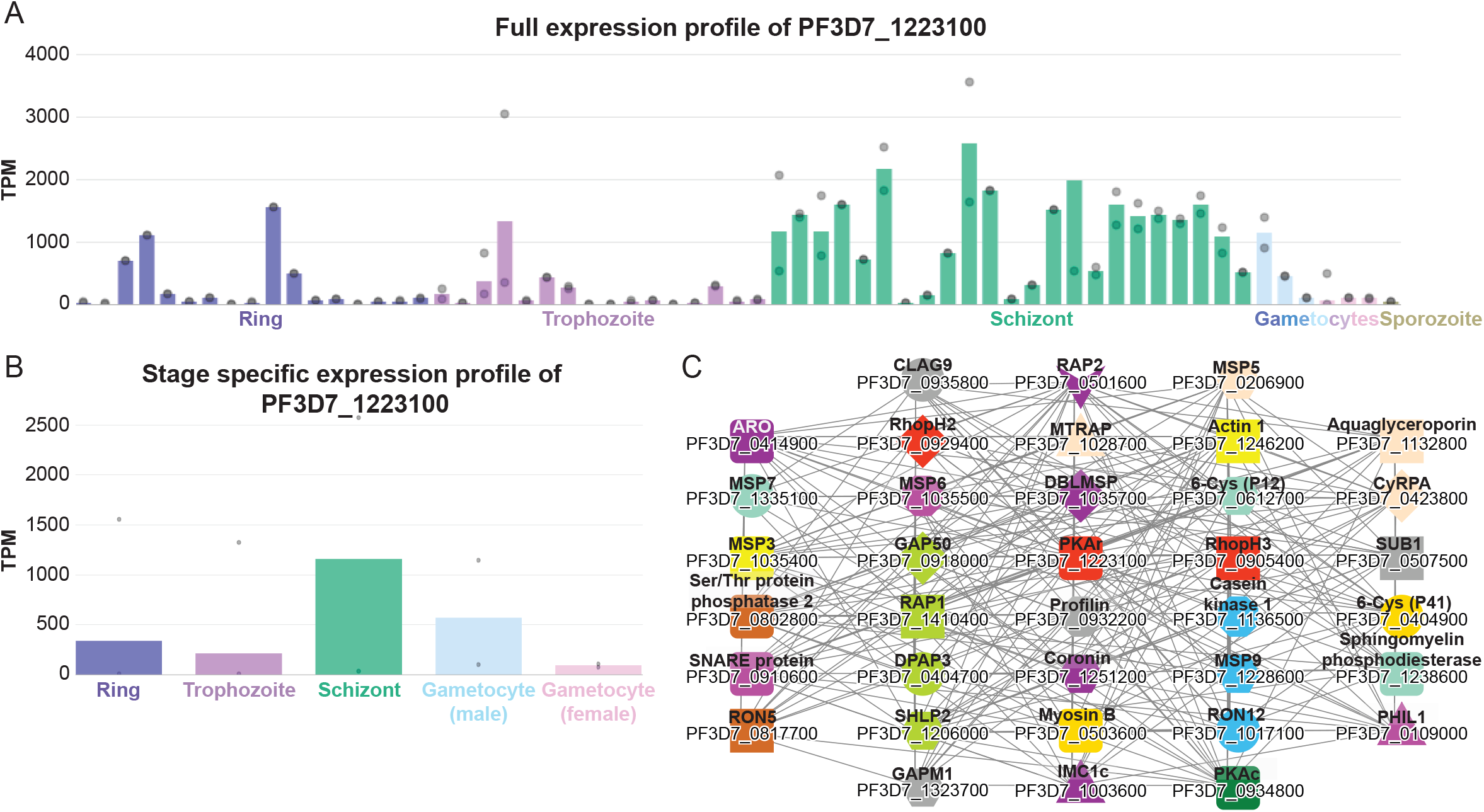
Expression profiles and co-expression neighborhood of gene *PF3D7_1223100*. A) Full expression profile of the gene. The x-axis represents the different RNA-seq experiments capturing the life stages and genetic perturbations of *P. falciparum*, while the y-axis indicates the expression level (Transcripts Per Million, TPM). The different life stages are color-coded by blue (ring), purple (trophozoite), green (schizont), light blue (male gametocyte), pink (female gametocyte) and brown (sporozoite). The bars indicate the mean expression value, while the dots show the expression values of the individual samples. For brevity, only the general descriptions of the samples are shown. B) Simplified expression profile of *PF3D7_1223100*, showing the average expression in the five major life stages. C) Co-expression network neighborhood containing functionally characterized genes. Nodes represent genes, edges (lines) connect co-expressed genes, while colored shapes indicate orthogroups. For brevity, only genes with experimentally verified function supported by at least two publications are shown in the figure.

### Comparative transcriptomic analysis of gene modules involved in male gametocyte-specific motility

Comparative transcriptomic analyses can reveal which gene modules are conserved across species (28, 29), thus enabling the identification of the core genetic components of specific biological processes (29, 30). Malaria.tools provides two methods to extract these conserved transcriptional programs by (i) identifying common gene families that are specifically expressed in a particular life stage of two malaria species or by (ii) identifying conserved clusters of co-expressed genes.

Using the first method to identify conserved transcriptional programs, we navigated to ‘Tools\Compare specificities’, selected species *P. falciparum* and *P. berghei*, set condition ‘Gametocyte (male) for both species and clicked ‘Compare specificity’. The database first identified genes that are preferentially expressed in male gametocytes in both species (specificity measure (SPM) > 0.85) (31), which revealed that 187 gene families are expressed at this life stage in the two parasites (Figure 2A, Table S6). The table below the Venn diagram shows the identity and links to the 187 gene families, and clicking on the tree links of a gene family depicted the phylogenetic and expression relationships of the genes in the family. Not surprisingly, many of the gene families show conserved male gametocyte-specific expression in the two *Plasmodium* species, as exemplified by the phylogenetic tree of gene family OG0000502 (Figure 2B, https://malaria.sbs.ntu.edu.sg/tree/view/503), indicating that this family is male gametocyte-specific. Interestingly, we also observed cases where only one of the clades of the phylogenetic tree showed a male gametocyte-specific expression (Figure 2C, OG0000055, https://malaria.sbs.ntu.edu.sg/tree/view/56), suggesting that for this particular gene family an ancient gene duplication took place in the ancestor of the *Plasmodium* species followed by either a sub-functionalization or neo-functionalization of the genes. The list of the conserved male-specific genes and gene families provides a good starting point to dissect the genetic basis of male gametocyte-specific biological processes.

**Figure 2.**
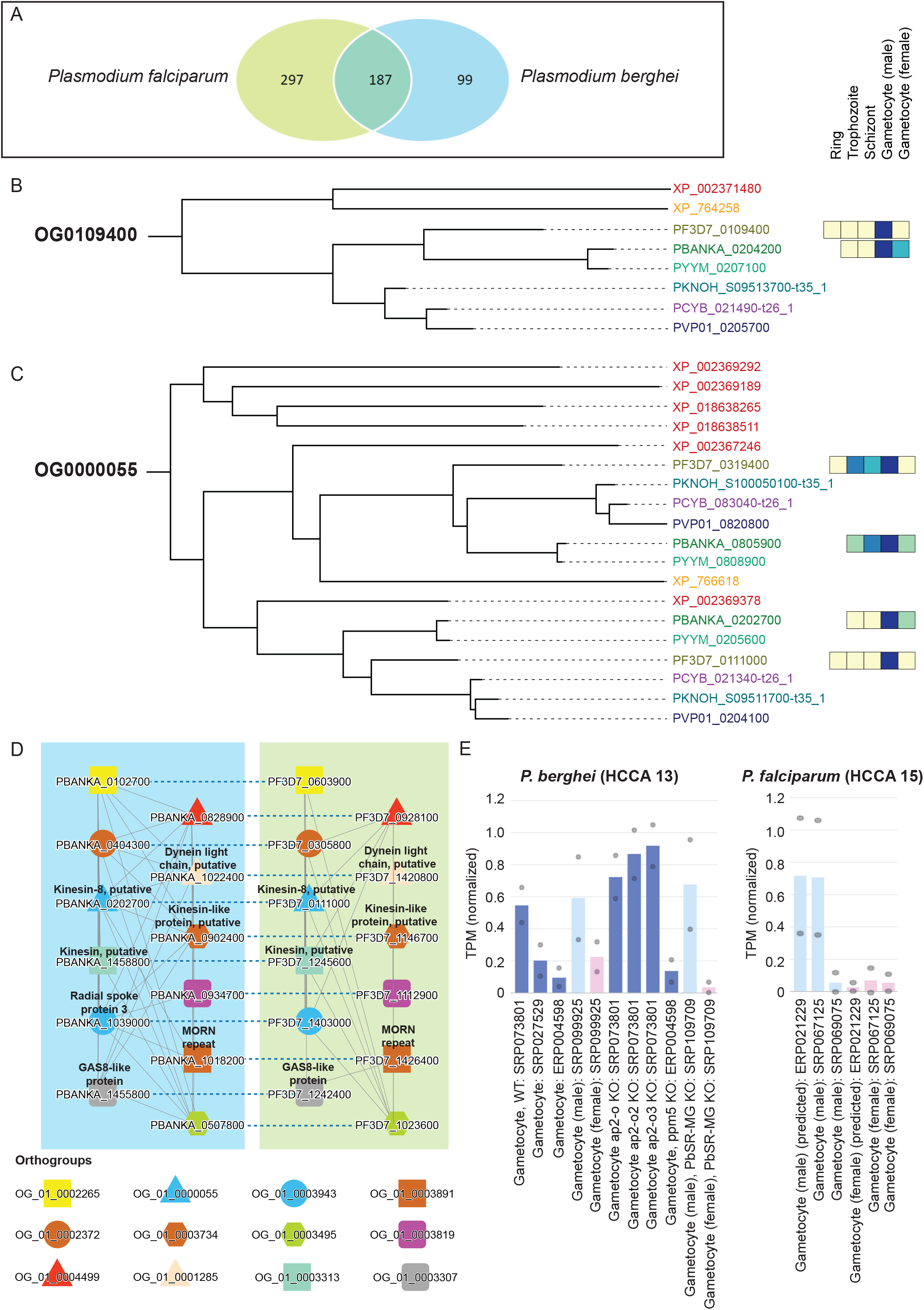
Comparative analysis of male gametocyte-specific gene expression in *P. berghei* and *P. falciparum*. A) Venn diagram showing the overlap of the male gametophyte-specific gene families in the two malaria species. B) Phylogenetic gene tree of gene family OG0109400. The different species are color coded and represent *Toxoplasma gondii* in red (gene IDs XP_NNNNNNNNN, NP_XXXXXX), *Theilera parva* in orange (gene IDs XP_NNNNNN), *P. falciparum* in olive (gene IDs PF3D7_XXXXXXX), *P. berghei* in dark green (gene IDs PBANKA_XXXXXXX), *P. yoleii* in light green (gene IDs PYYM_XXXXXXX), *P. knowlesi* in blue (gene IDs PKNOH_SXXXXXXXXX-t35_1), *P. vivax* in dark blue (gene IDs PVP01_XXXXXXX) and *P. cynomolgi* in purple (gene IDs PCYB_XXXXXX-t26_1). The colored boxes to the right of the gene IDs show the average gene expression in five major life stages of the malaria parasite, where yellow and blue color indicates low and high gene expression. C) Phylogenetic gene tree of gene family OG0000055. D) Comparison of cluster 13 (left blue box) and cluster 15 (right green box) from *P. berghei* and *P. falciparum*, respectively. Nodes represent, genes, solid edges connect co-expressed genes, dashed edges connect orthologs, while colored shapes indicate genes belonging to the same gene families. E) Average expression profiles of the genes found in cluster 13 (left) and 15 (right).

The second method to identify conserved gene modules using malaria.tools is based on co-expression network clusters. The clusters are used to identify functionally related genes based on the topology of the networks (9), i.e. similar clusters are found by identifying which cluster pairs contain significantly similar (P<0.05, hypergeometric test) number of gene families (expressed as Jaccard index,(5)). To exemplify this feature, we clicked first on one of the *P. berghei* genes found in the table (*PBANKA_0102700*, https://malaria.sbs.ntu.edu.sg/sequence/view/17914) and then on the co-expression cluster 13 containing this gene (https://malaria.sbs.ntu.edu.sg/cluster/view/40). The ‘Similar clusters’ table found on this page identified *P. falciparum* cluster 15 as being significantly similar to cluster 13 (Jaccard index = 0.246). Clicking on the ‘Compare’ button revealed the co-expression networks of the two modules (Figure 2D). As expected, the two conserved clusters show male gametocyte-specific expression profiles (https://malaria.sbs.ntu.edu.sg/cluster/view/40, https://malaria.sbs.ntu.edu.sg/cluster/view/124), further reinforcing that the two clusters represent a *bona fide* conserved transcriptional program for male gametocyte-specific motility.

In summary, the analysis of conserved gene modules resulted in 132 genes present in the co-expression networks whereof 123 genes showed male gametocyte-specific expression. However, only 60 of these homologs were annotated and the remaining ones were conserved proteins of unknown function (4 genes) or conserved *Plasmodium* proteins of unknown function (59 genes, Table S7). A closer inspection of the homologs present in the clusters revealed motor proteins such as kinesin (*PBANKA_0202700*, *PF3D7_0111000*, *PBANKA_0902400*, *PF3D7_1146700*, *PBANKA_1458800* and *PF3D7_1245600*) and dynein (*PBANKA_1022400* and *PF3D7_1420800*) (32), as well as flagella-related proteins such as the radial spoke protein 3 (*PBANKA_1039000*), growth arrest protein (33) (*PBANKA_1455800* and *PF3D7_1242400*) and MORN repeat containing protein which localises near the flagellar basal body in male gametocytes (34) (*PBANKA_1018200* and *PF3D7_1426400*). Taken together, the functions of these genes suggest an overall motility and flagella-related function associated with the clusters. Hence, the unannotated genes in these homologous networks will be of prime interest for researchers interested in male gametocyte motility.

### Identification of microneme-specific co-expressed gene clusters

Micronemes are protein rich, secretory organelles important for host-cell invasion and gliding motility in parasitic Apicomplexans (35). Proteins are being discharged to facilitate entry of the parasites into red blood cells.

To gain insight into microneme biogenesis and function using malaria.tools, we entered GO:0020009 (GO term for microneme) to arrive at the page dedicated to microneme cellular component (https://malaria.sbs.ntu.edu.sg/go/view/12866). The page revealed 270 and 299 annotated microneme-associated genes in *P. berghei* and *P. falciparum*, respectively. Furthermore, the page contains information about Pfam domains (Prot_kinase_dom, VWF_A, MORN and others) and gene families (OG_01_0000012, OG_01_0000038 and others), which may also play a role in the microneme function. Furthermore, the database identified cluster 2 (https://malaria.sbs.ntu.edu.sg/cluster/view/31) and cluster 7 (https://malaria.sbs.ntu.edu.sg/cluster/view/33) from *P. berghei* and cluster 2 (https://malaria.sbs.ntu.edu.sg/cluster/view/74) and cluster 12 (https://malaria.sbs.ntu.edu.sg/cluster/view/146) from *P. falciparum* as being significantly similar (P<0.05, Benjamini-Hochenberg corrected p-value)(36), implicating these clusters in a microneme-specific process.

To learn more about the function of these four clusters, we investigated their expression profiles. While cluster 2 from *P. berghei* and cluster 12 from *P. falciparum* show ubiquitous expression at all life stages of malaria (Figure 3A), cluster 7 from *P.berghei* and cluster 2 from *P. falciparum* show ookinete-and sporozoite-specific expression respectively. Since ookinetes and sporozoites are mosquito stage-specific, we speculate that the two *Plasmodium* species have at least two types of micronemes, one being ubiquitously expressed (clusters 2 and 12) and another being mosquito-specific (clusters 7 and 2).

**Figure 3.**
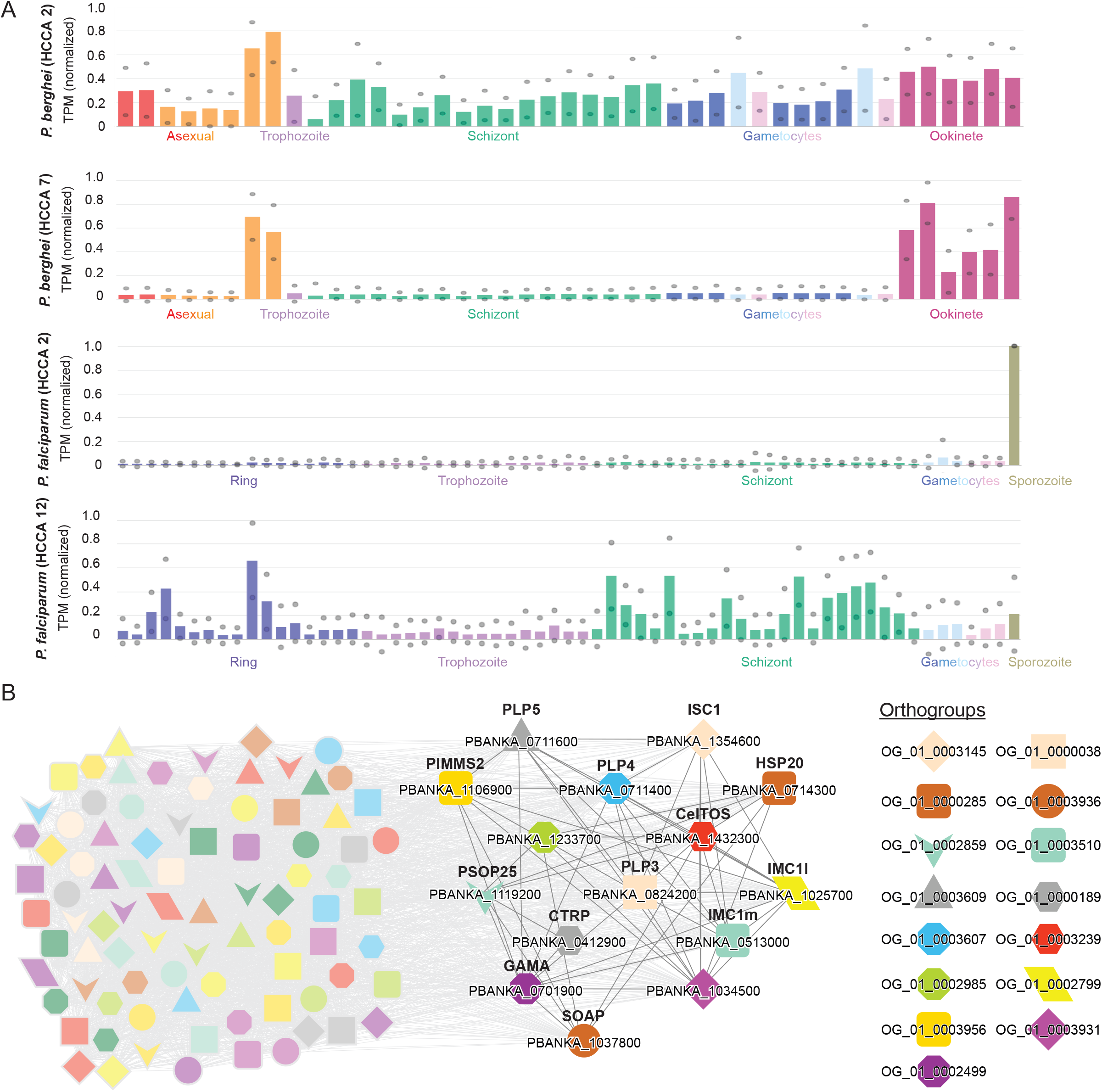
Expression profiles and co-expression network of the ookinete-enriched clusters in *P. falciparum* and *P. berghei*. A) Expression profiles of clusters 2 (first) and 7 (second) from *P. berghei* and clusters 2 (third) and 12 (fourth) from *P. falciparum*. The different life stages are color-coded. For brevity, the sample annotations are abbreviated to the major life stages. B) Co-expression cluster 7 from *P. berghei.* Nodes represent genes, co-expressed genes are connected by gray edges, while colored shapes indicate orthogroups. For brevity, only the discussed genes are highlighted.

We further investigated the putative function of the ookinete-specific cluster 7 from *P. berghei* (https://malaria.sbs.ntu.edu.sg/cluster/view/33), and found that the cluster is significantly enriched for GO terms such as entry into host cell (GO:0030260, Table S8), which is in line with the microneme being involved in invasion of blood cells. A closer look at the genes found in cluster 7 from *P. berghei* revealed that it contains genes that are essential for the infectivity and maturation of ookinetes (Figure 3B). Specifically, a group of genes relating to the inner membrane complex, surface ookinete protein and secreted ookinete protein is required for efficient gliding motility and midgut traversal [*PBANKA_1354600* (ISC1), *PBANKA_1025700* (IMC1l), *PBANKA_0513000* (IMC1m) (37), *PBANKA_1106900* (PIMMS2) (38), *PBANKA_0714300* (HSP20) (39) and *PBANKA_1432300* (CeITOS) (40)]. Another group of co-expressed genes contains perforins and a secreted protein, which are known to be important in midgut invasion where the parasite disrupts the membrane of the endothelial cell [*PBANKA_0824200* (PLP3), *PBANKA_0711400* (PLP4), *PBANKA_0711600* (PLP5) (41) and *PBANKA_1037800* (SOAP) (42)]. Last but not least, genes important for the transition from the ookinete to oocyst stage are also found enriched in this cluster [*PBANKA_0701900* (GAMA/PSOP9) (43), *PBANKA_1119200* (PSOP25) (44) and *PBANKA_0412900* (CTRP) (45)]. Taken together, cluster 7 from *P. berghei* contain genes that are most likely important for microneme function and host cell invasion.

## CONCLUSIONS

The lack of comparative genomic and transcriptomic resources for malaria prompted us to construct malaria.tools, a state-of-the-art database containing a wide range of user-friendly features. The database can be queried by BLAST, gene IDs, keywords and Pfam domains and Gene Ontology searches. To identify novel genes relevant for a biological process of interest, the co-expression neighborhoods and clusters can be mined for uncharacterized candidates that are connected to well-studied genes. Alternatively, the database allows an easy identification of genes that are expressed during a specific life stage of the malaria parasite, thus allowing researchers to dissect the transcriptome critical for pathogenicity and other life stages. Finally, the database can compare the clusters and stage-specific expression profiles to identify the conserved core components of various biological processes. We envision that malaria.tools will aid malaria researchers in selecting relevant genes for experimental functional characterisation and potential drug development for successful combating the emerging drug resistances.

## DATA AVAILABILITY

The expression matrices, RNAseq sample annotation and gene families are available from the supplementary material. The co-expression networks, coding and protein sequences can be downloaded from malaria.tools.

## SUPPLEMENTARY DATA

The Supplementary Data are available at NAR Online.

## ACKNOWLEDGEMENTS

Malaria.tools is hosted at Nanyang Technological University Singapore and we would like to thank Ryan Chee Kiang Ng for excellent tech support. We would like to thank Dr. Daniela Mutwil-Anderwald for proofreading the manuscript. Furthermore, we would like to thank Dr. Lei Zhu from Prof. Zbynek Bozdech lab, SBS, NTU for useful discussions.

## Author Contributions

Malaria.tools was implemented by Q.W.T. who also prepared the data and built malaria.tools with input from M.M. Both Q.W.T. and M.M. wrote the manuscript.

---

Table S1. TPM expression matrix for *Plasmodium berghei*. Genes are found in rows, while samples are found in columns.

Table S2. TPM expression matrix for *Plasmodium falciparum*. Genes are found in rows, while samples are found in columns.

Table S3. Annotation of RNAseq samples for *Plasmodium berghei*.

Table S4. Annotation of RNAseq samples for *Plasmodium falciparum*.

Table S5. Orthogroups of the eight Apicomplexa species included in this study. Each row represents one orthogroup and the genes belonging to the orthogroup.

Table S6. Male gametocyte specific genes of *Plasmodium falciparum* and *Plasmodium berghei*.

Table S7. Annotation of comparative clusters *Plasmodium berghei* cluster 13 and *Plasmodium falciparum* cluster 15

Table S8. Enriched GO terms of *P. berghei* HCCA cluster 7 and genes associated with it.

**Figure S1.**
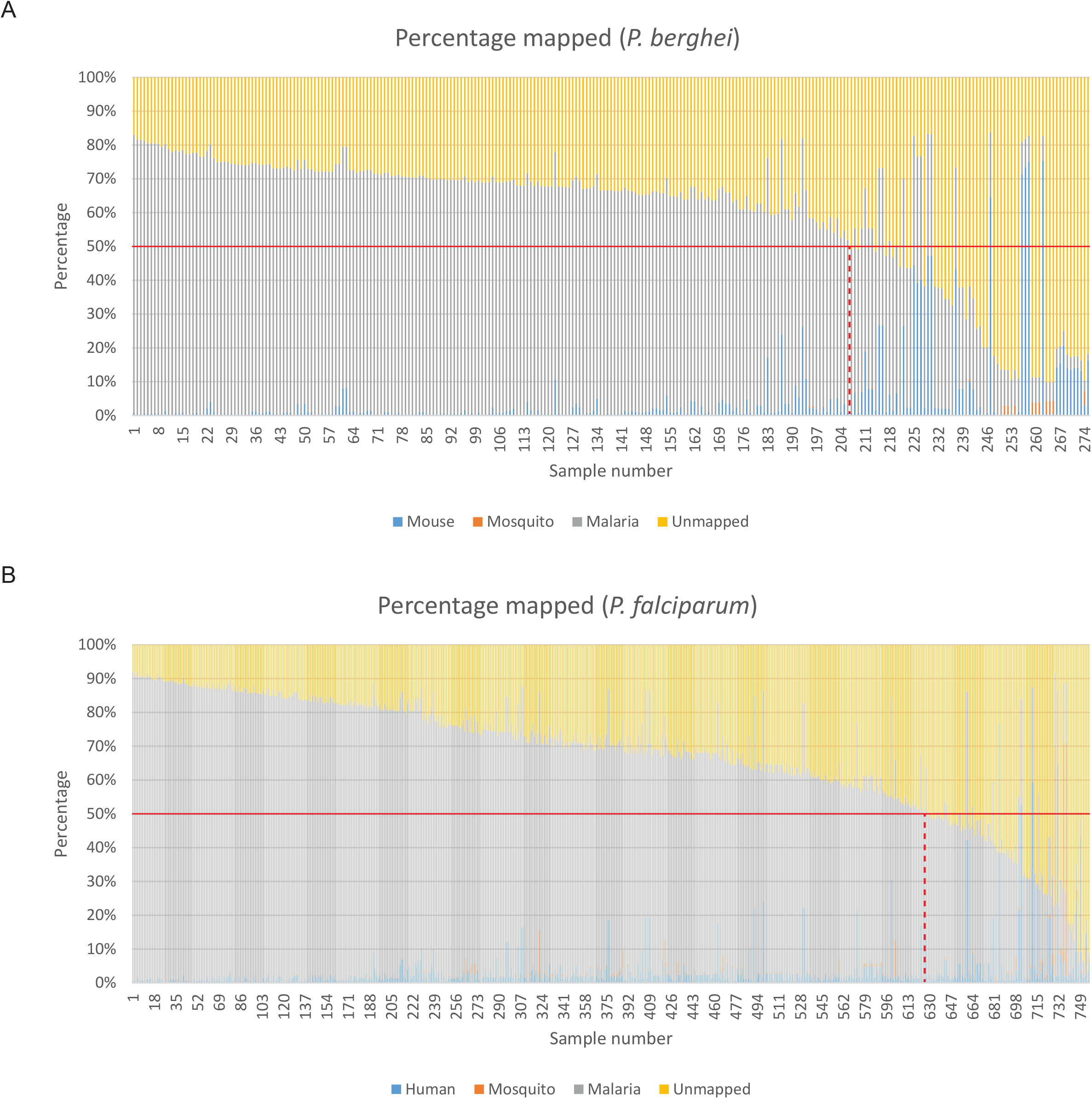
Overview of the sample quality of the RNA-seq samples that contain at least 1 million reads. A) *Plasmodium berghei*. For each sample, the percentage of reads mapping to the parasite (grey), mouse or human (blue), mosquito (orange) and not mapping to either (yellow) is shown. The sample cutoff is indicated by the red line. B) Overview of the sample quality of *Plasmodium falciparum*.

